# Fatty acids produced by the gut microbiota dampen host inflammatory responses by modulating intestinal SUMOylation

**DOI:** 10.1101/2022.01.09.475538

**Authors:** Chaima Ezzine, Léa Loison, Christine Bole-Feysot, Pierre Déchelotte, Moïse Coëffier, David Ribet

## Abstract

The gut microbiota produces a wide variety of metabolites, which interact with intestinal cells and contribute to host physiology. These metabolites regulate intestinal cell activities by modulating either gene transcription or post-translational modifications of gut proteins. The effect of gut commensal bacteria on SUMOylation, an essential ubiquitin-like modification in intestinal physiology, remains however unknown. Here, we show that short chain fatty acids (SCFAs) and branched chain fatty acids (BCFAs) produced by the gut microbiota increase protein SUMOylation in different intestinal cell lines in a pH-dependent manner. We demonstrate that these metabolites induce an oxidative stress which inactivates intestinal deSUMOylases and promotes the hyperSUMOylation of chromatin-bound proteins. In order to determine the impact of these modifications on intestinal physiology, we focused on the NF-κB signaling pathway, a key player in inflammation known to be regulated by SUMOylation. We demonstrated that the hyperSUMOylation induced by SCFAs/BCFAs inhibits the activation of the NF-κB pathway in intestinal cells by blocking the degradation of the inhibitory factor IκBα in response to TNFα. This results in a decrease in pro-inflammatory cytokines expression, such as IL8 or CCL20, as well as a decrease in intestinal epithelial permeability in response to TNFα. Together, our results reveal that fatty acids produced by gut commensal bacteria regulate intestinal physiology by modulating SUMOylation and illustrate a new mechanism of dampening of host inflammatory responses by the gut microbiota.

## Introduction

The gut microbiota produces a wide variety of metabolites diffusing to the intestinal mucosa and modulating intestinal cell activities^1^. Some of these metabolites may even cross the intestinal barrier and reach distant organs via the bloodstream or via nerve communications. Fatty acids constitute a major class of metabolites produced by intestinal bacteria. They include the so-called Short Chain Fatty Acids (SCFAs), which are carboxylic acids with aliphatic tails of 1 to 6 carbons^2^. Acetate, butyrate and propionate are the main SCFAs produced in the human colon and derive from the anaerobic catabolism of dietary fibers and proteins by intestinal bacteria^3,4^. Branched Chain Fatty Acids (BCFAs), such as isobutyrate, isovalerate or 2-methylbutyrate, constitute another class of fatty acids produced by bacteria with one or more methyl branches on the carbon chain. BCFA mostly derive from the breakdown of proteins by intestinal bacteria, and more particularly from the catabolism of branched-chain amino-acids (valine, leucine and isoleucine, producing isobutyrate, isovalerate or 2-methylbutyrate, respectively)^5^.

Fatty acids regulate intestinal cell activities by various mechanisms. They may bind to specific receptors expressed on intestinal cells, such as GPR41/FFAR3, GPR43/FFAR2 and GPR109A, and activate various signaling pathways^6^. Fatty acids may also directly enter into intestinal cells by passive diffusion or by facilitated transport. Once in intestinal cells, they participate to the cell metabolism. For example, colonocytes were shown to use butyrate as a major energy source or, alternatively, isobutyrate when butyrate availability is low^7,8^. Finally, fatty acids may regulate intestinal cell activities by interfering with post-translational modification such as neddylation^9,10^. The impact of fatty acids on other ubiquitin-like modifications in intestinal cells has not been described yet.

SUMOylation is an ubiquitin-like modifications consisting in the covalent addition of SUMO (<u>S</u>mall <u>U</u>biquitin-like <u>MO</u>difier) peptides to target proteins. Five SUMO paralogs have been identified in humans that share 45-97% sequence identity. SUMO1, SUMO2 and SUMO3, which are the most studied paralogs, can be conjugated to both overlapping and distinct sets of proteins^11^. The conjugation of SUMO to lysine residues of target proteins is catalysed by an enzymatic machinery composed of one E1 enzyme (SAE1/SAE2), one E2 enzyme (UBC9) and several E3 enzymes^12^. SUMOylation is a reversible modification as the isopeptide bond between SUMO and its target can be cleaved by specific proteases called deSUMOylases^13^. The consequences of SUMO conjugation on target proteins are very diverse and include changes in protein localization, stability, activity or interactions with other cellular components^11,14,15^.

SUMOylation plays essential roles in intestinal physiology as it limits detrimental inflammation while participating to tissue integrity maintenance^16,17^. Interestingly, several intestinal bacterial pathogens were shown to interfere with epithelial cell SUMOylation^18^. *Listeria monocytogenes*, for example, secretes a pore-forming toxin triggering the degradation of the host cell E2 SUMO enzyme and the rapid loss of SUMO-conjugated proteins^19,20^. *Salmonella enterica* serovar Typhimurium also targets the host E2 SUMO enzymes during infection by inhibiting its translation via miRNA-based mechanisms^21^. *Shigella flexneri*, finally, similarly switches off the SUMOylation machinery by triggering a calpain-dependent cleavage of the SUMO E1 enzyme SAE2 in infected cells^22^. In contrast to these examples of pathogens dampening intestinal cell SUMOylation, the impact of gut commensal bacteria on the SUMOylation of intestinal proteins remains unknown. We investigate here whether bacterial metabolites derived from the gut microbiota regulate intestinal cell activities by modulating host protein SUMOylation. We demonstrate that bacterial fatty acids induce an hyperSUMOylation in intestinal cells, which dampens inflammatory responses and promotes intestinal epithelial integrity.

## Material and Methods

### Animals

Animal care and experimentation were approved by a regional Animal Experimentation Ethics Committee (APAFIS#21102–2019061810387832 v2) and complied with the guidelines of the European Commission for the handling of laboratory animals (Directive 2010/63/EU). All efforts were made to minimize suffering of animals.

Eight-weeks-old C57Bl/6JRj male mice (Janvier Labs, Le-Genest-Saint-Isle, France) were housed at 23°C (5 animals/cage) with a 12-h light-dark cycle in regular open cages. All animals were fed with a non-sterilized standard rodent diet (3430.PM.S10, Serlab, France). Drinking water was not sterilized. After 1 week of acclimatization to the animal facility, animals were split in two groups (5-10 animals/group): one group received antibiotics by oral gavage once a day, while the other group had no antibiotic treatment and were gavaged once a day with drinking water. For oral gavages, mice received a volume of 10 _μ_L/g body weight of drinking water supplemented with 0.1 mg/mL Amphotericin-B (Simag-Aldrich), 10 mg/mL Ampicillin (Sigma-Aldrich), 10 mg/mL Neomycin trisulfate salt hydrate (Simag-Aldrich), 10 mg/mL Metronidazole (Simag-Aldrich) and 5 mg/mL Vancomycin hydrochloride (Simag-Aldrich)^23^. This solution was delivered with a stainless steel tube without prior sedation of the mice. To prevent fungal overgrowth in the antibiotic-treated animals, mice were pre-treated with Amphotericin-B for 3 days before the beginning of the protocol^23^. As for antibiotic treatment, Amphotericin-B was delivered by oral gavage (10 μL/g body weight of drinking water supplemented with 0.1 mg/mL Amphotericin-B)^23^. Three independent animal series were performed. At the end of the study, all animals were euthanized by an intraperitoneal injection of an overdose of ketamine (200 mg/kg BW) and xylazine (20 mg/kg BW). Jejunal and caecal segments, as well as cecal content were then removed, frozen in liquid nitrogen and stored at −80°C. Two independent animal series were performed.

### Quantification of caecal microorganisms by quantitative PCR

Quantitative real-time polymerase chain reaction (qPCR) was performed on DNA samples extracted from mice cecal contents to monitor the efficiency of bacterial depletion in mice treated with antibiotics, as described in ref. 23. To quantify total Eubacteria, qPCR were performed using primers targeting the bacterial 16S rRNA gene (Eub-338F, 5’-ACTCCTACGGGAGGCAGCAG-3’ and Eub-518R, 5’-ATTACCGCGGCTGCTGG-3’)^24^. The Cq determined in each sample were compared with a standard curve made by diluting genomic DNA extracted from a pure culture of *E. coli*, for which cell counts were determined prior to DNA isolation.

### Protein extraction from mouse intestinal tissues

Intestinal tissues were mechanically lysed using bead beating in a buffer containing 50 mM HEPES pH 8.0, 8 M urea buffer, supplemented with 10 mM N-ethyl-maleimide (NEM). Tissue lysates were then centrifugated for 15 min at 13,000x*g* at 4°C. Supernatents were collected, mixed with one volume of Laemmli buffer (125 mm Tris-HCl [pH 6.8], 4% SDS, 20% glycerol, 100 mm dithiothreitol [DTT], 0.02% bromphenol blue) and anlyzed by immunoblotting.

### Cell culture

CACO2 (American Type Culture Collection (ATCC)-HTB-37), HeLa (ATCC-CCL2) and T84 (ATCC CCL-248) cells were cultivated at 37°C in a 5% CO_2_ atmosphere. CACO2 and HeLa cells were cultivated in Minimum Essential Medium (MEM) (Eurobio) supplemented with 2 mM L-Glutamine (Invitrogen), 10% Fetal Bovine Serum (FBS, Eurobio), non-essential aminoacids (Sigma-Aldrich), 1 mM sodium pyruvate (Gibco) and a mixture of penicillin (10000U/mL) and streptomycin (10mg/mL). T84 cells were cultivated in DMEM/F12 (Dulbecco’s Modified Eagle Medium F-12) (Eurobio) supplemented with 10% FBS and 2.5 mM L-Glutamine.

CACO2 and T84 cells were seeded in wells at a density of 1.1×10^5^ cells/cm^2^ and 1.7×10^5^ cells/cm^2^, respectively, the day before treatments with BCFA or SCFA.

Before treatments, cell culture medium was replaced by HBSS (Hanks’ Balanced Salt Solution; Sigma-Aldrich). Cells were then treated as indicated in the text. For BCFAs and SCFAs treatments, 100 mM stock solutions in water were first prepared from the corresponding acidic form (*e*.*g*. isobutyric acid) or from the sodium salt of the corresponding basic form (*e*.*g*. sodium isobutyrate) and then further diluted in cell culture media (HBSS). When needed, the pH of cell culture medium was shifted using either 0.1 M NaOH or 0.1 M HCl solution. For treatments with ROS inhibitors, CACO2 cells were pre-incubated for 30 min with 5 mM N-acetyl-cysteine (NAC) or 10 µM Diphenyleneiodonium (DPI) and then incubated for 1 h with 5 mM isobutyric acid or isovaleric acid. For immunoblotting experiments, cells were lysed directly in Laemmli buffer. For TNFα treatments, CACO2 cells were first incubated with BCFAs or SCFAs for 1 hour and then incubated with 100 ng/mL TNFα. For immunoblotting and qRT-PCR analyses, cells were lysed after 30 min or 1h of incubation with TNFα, respectively. For Transepithelial electrical resistance (TEER) measurements, cells were incubated for 24h with TNFα.

### Immunoblot analyses

Cell lysates and protein extracts from intestinal tissues in Laemmli buffer were boiled for 5 min, sonicated and protein content was analysed by electrophoresis on TGX Stain-free pre-cast SDS-polyacrylamide gel (Bio-rad). Proteins were then transferred on PVDF membranes (GE Healthcare) and detected after incubation with specific antibodies using ECL Clarity Western blotting Substrate (Bio-Rad). Primary and secondary antibodies used for immunoblot analyses are described in Supplementary Table S1. All displayed immunoblots are representative of at least two independent experiments. Quantifications of proteins from intestinal tissues were performed on a ChemiDoc Imaging System (Bio-rad). SUMO2/3-conjugated proteins levels (above 50 kDa) were normalized by the level of total proteins in each lysate (determined using the TGX-stain free imaging technology; Bio-rad).

### Detection of Reactive Oxygen Species

Detection of ROS was adapted from ref. 25. Luminol was dissolved in NaOH 0.1 M to obtain a 50 mM stock solution. A stock solution of 1000 U/mL HRP (HorseRadish Peroxidase) was prepared in parallel in PBS (Phosphate-Buffered Saline). Culture media from CACO2 and HeLa cells treated with BCFAs or SCFAs were collected and centrifugated for 5 min at 13,000x*g* at room temperature to eliminate cell remnants. The pH of the obtained supernatents was then buffered to 7.5 to avoid pH-dependent interferences with luminol activity. Luminol (1 mM final concentration) and HRP (4 U/mL) were finally added to each culture media and luminescence was quantified immediately on a luminometer (Tecan).

### DeSUMOylase activity

DeSUMOylase activity assays were adapted from ref. 26. CACO2 and T84 cells grown in 12-well plates were scraped in 100 µL lysis buffer (Tris HCl pH 8.0 50 mM, EDTA 5 mM, NaCl 200 mM, Glycerol 10%, NP40 0.5%). Negative controls were prepared by adding 10 mM N-ethymaleimide (NEM; Sigma-Aldrich) to cell lysates. Recombinant human SUMO1-AMC and SUMO2-AMC proteins (R&D Systems) were diluted in parallel to 500 nM in Assay buffer (Tris HCl pH 8.0 50 mM, Bovine Serum Albumin (BSA) 100 µg/mL, Dithiothreitol (DTT) 10 mM). For each measurement, 10 µL of cell lysate were mixed with 40 µL of SUMO-AMC containing Assay buffer and fluorescence (λ_Ex_=380 nm; λ_Em_=460 nm) was recorded for 30 min at 37°C on a Flexstation 3 microplate reader (Molecular Devices). Measurements were performed in duplicate. DeSUMOylase activities were determined by calculating the initial speed of fluorescence emission in each lysate, normalized by the quantity of proteins in the corresponding sample, determined in parallel using BCA assays (Pierce™ BCA Protein Assay Kit).

### Cell fractionation

CACO2 cells incubated or not with 5 mM isobutyric, isovaleric or butyric acids for 5 h were washed once at room temperature in PBS, collected by scraping in ice-cold PBS, and then centrifugated at 130x*g* at 4°C for 3 min. Cell pellets were then lysed in 5 volumes of ice-cold E1 buffer (50 mM HEPES-KOH pH 7.5; 140 mM NaCl; 1mM EDTA; 10% glycerol; 0.5% NP-40; 0.25% Triton X-100; 1 mM DTT; protease inhibitors [complete protease inhibitor cocktail tablets; Roche]) and centrifugated at 1,100x*g* at 4°C for 2 min. The supernatants (corresponding to cytosolic fractions) were then collected. Pellets were washed by adding 5 volumes of E1 buffer and centrifugated at 1,100x*g* at 4°C for 2 min. Pellets were additionally washed by adding 5 volumes of E1 buffer, incubated 10 min on ice and centrifugated at 1,100x*g* at 4°C for 2 min. Pellets were then resuspended in 2 volumes of ice-cold E2 buffer (10 mM Tris-HCl pH 8.0; 200 mM NaCl; 1 mM EDTA; 0.5 mM EGTA; protease inhibitors). These suspensions were centrifugated at 1,100xg at 4°C for 2min and supernatants (corresponding to nuclear soluble fractions) were collected. Pellets were washed by adding 2 volumes of E2 buffer and centrifugated at 1,100x*g* at 4°C for 2 min. Pellets were additionally washed by adding 2 volumes of E2 buffer, incubated 10 min on ice and centrifugated at 1,100x*g* at 4°C for 2 min. Pellets were then resuspended in 5 volumes of ice-cold E3 buffer (500 mM Tris-HCl pH 6.8; 500 mM NaCl; protease inhibitors). These suspensions were centrifugated at 16,000x*g* at 4°C for 10 min. Pellets (corresponding to insoluble chromatin-bound fractions) were finally resuspended directly in Laemmli buffer. Cytosolic and nuclear soluble fractions were mixed 1:1 with Laemmli buffer. All fractions were boiled for 5 min and sonicated before immunoblotting analyses.

### Quantification of proinflammatory cytokines expression

Total RNAs were extracted from CACO2 cells using RNeasy Plus Mini kit (Qiagen) following manufacturer’s instructions. For each condition, 1 µg of total RNAs was reverse transcribed using random hexamers and M-MLV reverse transcriptase (Invitrogen). Specific cDNAs were then quantified by qPCR using Itaq Universal SYBR Green Supermix (Bio-Rad). GAPDH was used as an internal reference for normalization. Primers used in this study are hGAPDH_F (5’-TGCCATCAATGACCCCTTCA-3’), hGAPDH_R (5’-TGACCTTGCCCACAGCCTTG-3’), hIL8_F (5’-TGGCAGCCTTCCTGATTT-3’), hIL8_R (5’-AACTTCTCCACAACCCTCTG-3’), hCCL20_F (5’-TTTGCTCCTGGCTGCTTTGA-3’) and hCCL20_R (5’-GCAAGTGAAACCTCCAACCC-3’). Serial dilution of target cDNAs were included on each plate to generate a relative curve and to integrate primer efficiency in the calculations of mRNA quantities.

### Evaluation of intestinal epithelial permeability

CACO2 cells were seeded in Transwell inserts and cultivated for 21 days. Monolayer formation and differenciation was monitored by daily evaluation of transepithelial electrical resistance (TEER) measurement, performed with an EVOM epithelial voltohm meter equipped with ‘‘chopstick’’ electrodes. After three weeks, cell culture media were replaced by HBSS. Cells were preincubated or not with isobutyric or isovaleric acids for 1 hour. 100 ng/mL TNFα was then added to both apical and basolateral compartments. TEER was evaluated after 24h of incubation.

## Results

### Gut microbiota depletion decreases protein SUMOylation in the caecum

To determine whether the gut microbiota affects intestinal SUMOylation, we compared the global SUMOylation patterns of intestinal segments from conventional mice or from mice with a depleted gut microbiota. Depletion of mice intestinal bacteria was performed via the oral gavage of a cocktail of antibiotics during 7 days^23^. We quantified by qPCR the total amount of Eubacteria in the cecal content of mice treated with antibiotics to ensure that the efficiency of bacterial depletion was above 75%. The SUMOylation pattern of jejunal and caecal segments were then analyzed by immunoblotting experiments using anti-SUMO2/3 antibodies (Fig. 1). The level of SUMO2/3-conjugated proteins (above 50 kDa) was quantified in each sample. Interestingly, we observed that mice with a depleted gut microbiota exhibit a significant decrease in the level of SUMO2/3-conjugated proteins in the caecum (Fig. 1). In contrast, we did not observe any significant modification of the SUMOylation profile in the jejunum of mice treated with antibiotics (Fig. 1). Together, these results suggest that the gut microbiota regulates the level of protein SUMOylation in specific intestinal segments.

**Figure 1:**
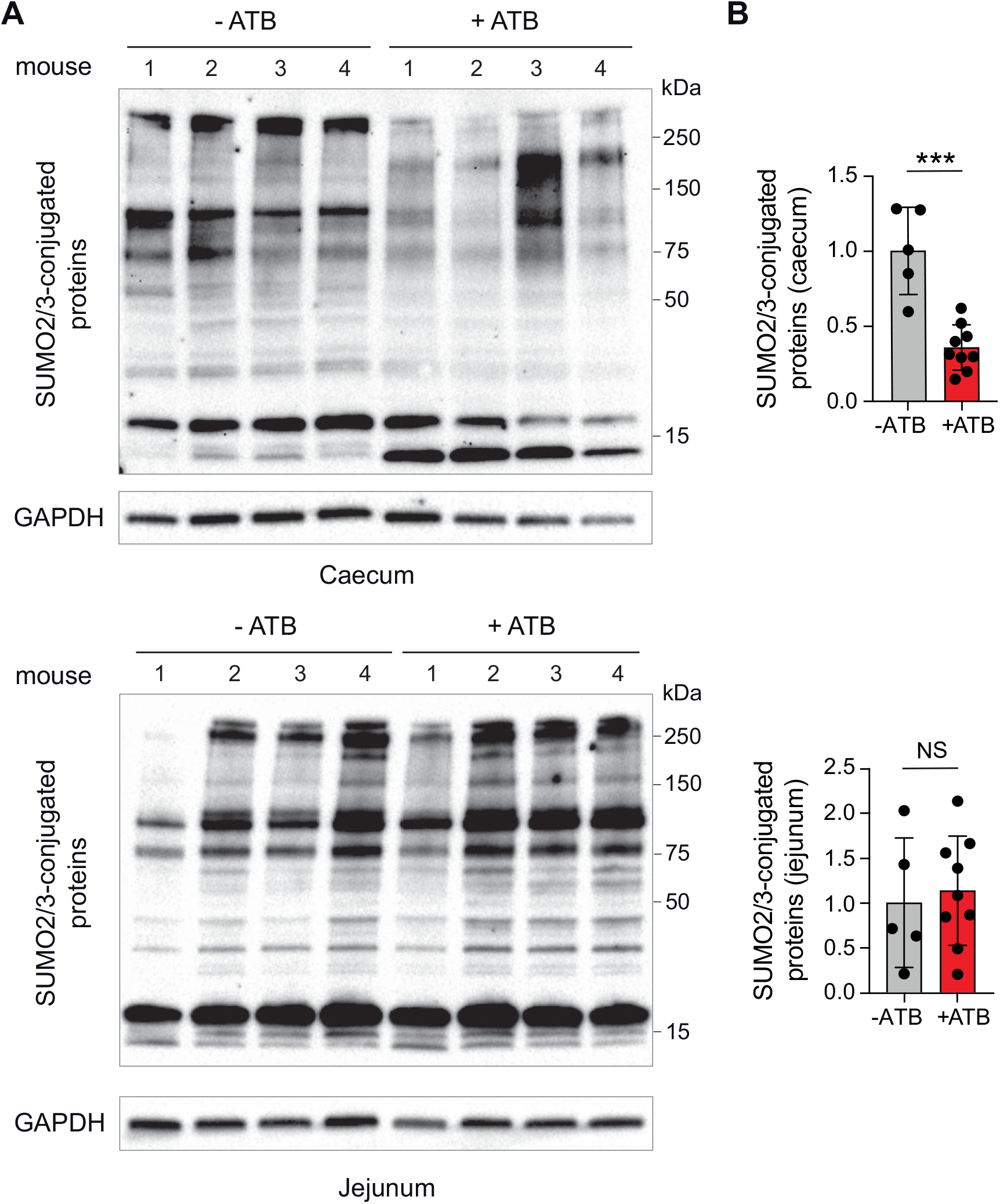
Gut microbiota depletion decreases protein SUMOylation in the caecum. A, Immunoblot analysis of SUMO2/3-conjugated proteins and GAPDH levels in the caecum and jejunum of mice treated or not with antibiotics (ATB) (4 representative mice are shown for each group). B, Quantification of SUMO2/3-conjugated proteins. Values are expressed as fold-change versus untreated mice (mean ± s.d.; *n*=5-9; ***, *P*<0.001; NS, not significant; two-tailed Student’s t-test).

### BCFAs trigger an hyperSUMOylation of intestinal proteins

As fatty acids such as SCFAs and BCFAs are important mediators of the interactions between gut bacteria and host cells, we assessed if these metabolites may modulate intestinal cell SUMOylation. We first monitored the effect of BCFAs on intestinal cell SUMOylation *in vitro* by incubating CACO2 cells for 1h or 5h with isobutyric, isovaleric or 2-methyl-butyric acids (1 mM or 5 mM final concentrations) (Fig. 2 and S1). Interestingly, we observed that all BCFAs induced an increase in the level of proteins conjugated to SUMO2/3 after 1h or 5h of incubation (at 5 mM concentration). We observed similar results in another intestinal cell line, T84 cells, after incubation with isobutyric, isovaleric acids or 2-methyl-butyric acids (Fig. 2 and S1).

**Figure 2:**
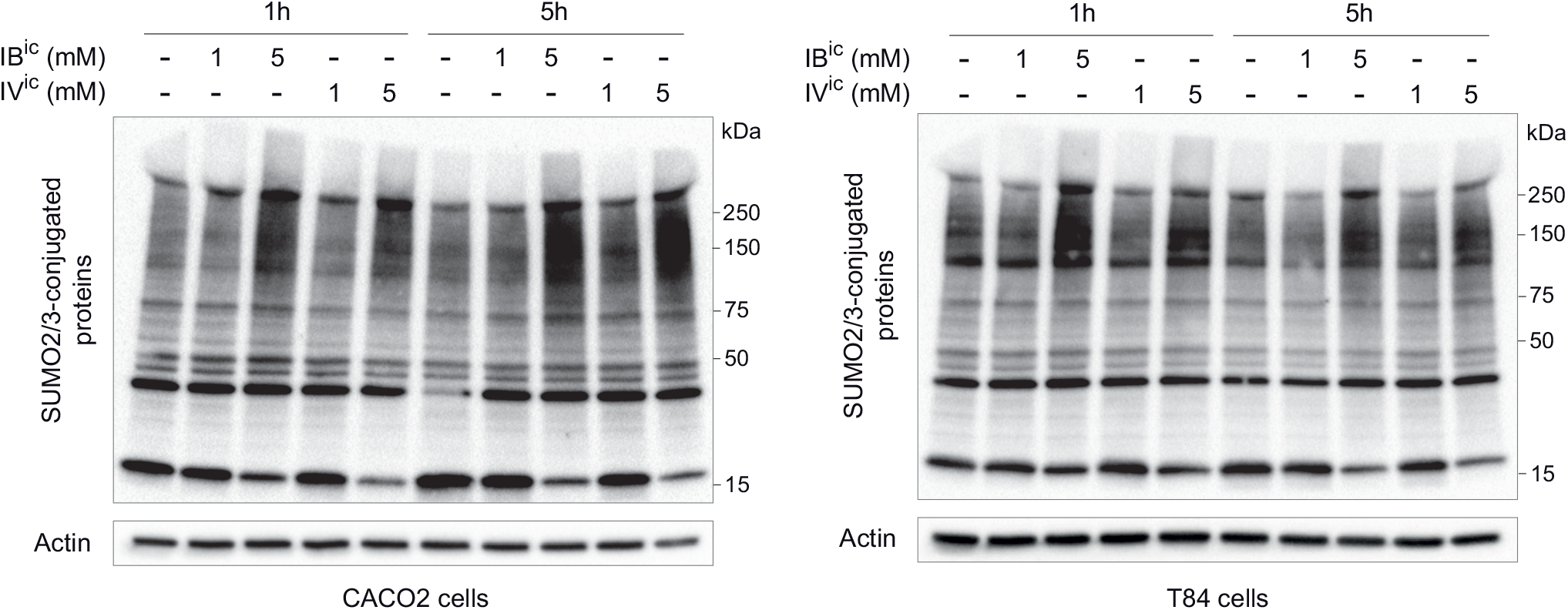
BCFAs trigger hyperSUMOylation of intestinal proteins *in vitro*. Immunoblot analysis of SUMO2/3-conjugated proteins and actin levels in CACO2 (left) and T84 (right) cells incubated with isobutyric acid (IB^ic^) or isovaleric acid (IV^ic^) for 1 or 5 h.

### The effect of BCFAs on intestinal SUMOylation is pH dependent

BCFAs are weak organic acids, which exist in solution either as acidic (R-COOH) or basic (R-COO^-^) forms. For example, addition of 5 mM isovaleric acid in HBSS medium leads to a solution with a pH of ∼5.2 containing approximatively 30% (*i*.*e*. ∼1.4 mM) of the acidic form (isovaleric acid) and 70% (*i*.*e*. ∼3.6 mM) of the basic form (isovalerate). In contrast, addition of 5 mM sodium isovalerate in HBSS medium leads to a solution with a pH of ∼7.5 containing approximatively 0.2% (*i*.*e*. ∼0.01 mM) of isovaleric acid and 99.8% (*i*.*e*. ∼4.99 mM) of isovalerate. To decipher whether both acidic and basic forms of BCFAs trigger hyperSUMOylation in intestinal cells, we added 5 mM isovaleric acid to CACO2 cells supernatant and increased gradually the cell culture medium pH from 5.2 to 7.0 (thereby decreasing the isovaleric acid/isovalerate ratio) (Fig. 3A). We did not observe any hyperSUMOylation when cells where incubated at pH 6.0 or 7.0, in contrast to cells incubated at pH 5.2. This shows that only isovaleric acid (and not isovalerate) promotes SUMO-conjugation of intestinal proteins.

**Figure 3:**
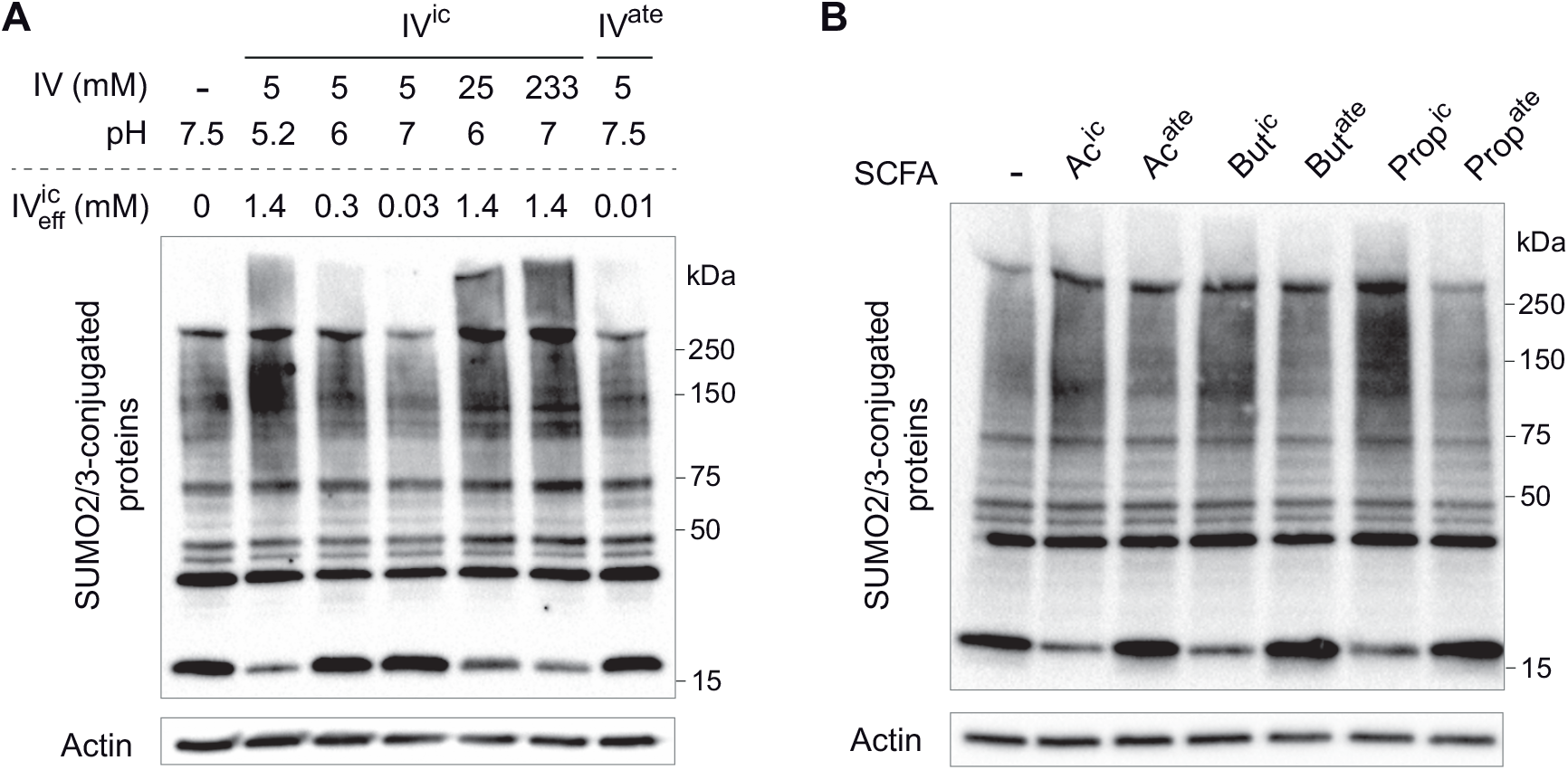
BCFAs and SCFAs trigger hyperSUMOylation of intestinal proteins in a pH-dependent manner. A, Immunoblot analysis of SUMO2/3-conjugated proteins and actin levels in CACO2 cells incubated with various concentrations of isovaleric acid (IV^ic^) or sodium isovalerate (IV^ate^) at definite pH. The pH and the corresponding effective concentrations of isovaleric acid (IV^ic^_eff_) is indicated for each condition. B, Immunoblot analysis of SUMO2/3-conjugated proteins and actin levels in CACO2 cells incubated for 5 h with 5 mM acetic acid (Ac^ic^), sodium acetate (Ac^ate^), butyric acid (But^ic^), sodium butyrate (But^ate^), propionic acid (Prop^ic^) or sodium propionate (Propate).

We then added increasing amounts of isovaleric acid to CACO2 cell culture medium and set in parallel the pH between 5.2 and 7.0. For each pH, the amount of isovaleric acid added to cells was calculated to maintain a final concentration of isovaleric acid in the cell culture medium to ∼1.4 mM. We observed that this increase in BCFA concentration restores the hyperSUMOylation observed in CACO2 cells at pH 6 and 7 (Fig. 3A). This result demonstrates that BCFAs, when present in high concentration, trigger hyperSUMOylation even at neutral or weakly acidic pH.

Of note, the acidic forms of fatty acids are uncharged and freely diffusible across cellular membranes, in contrast to the basic forms, which are negatively charged and only cross membranes thanks to specific transporters. Thus, as only the acidic forms of BCFA induce an hyperSUMOylation of intestinal proteins, we can hypothetise that these forms diffuse passively across the cell membrane, and then act intracellularly on intestinal cell SUMOylation.

### SCFAs also affect intestinal SUMOylation

To complete our results obtained with BCFAs, we determined whether SCFAs similarly impact intestinal cell SUMOylation. We incubated CACO2 cells with acetic, butyric and propionic acid for 5h (5 mM final concentration; pH∼5.2). Interestingly, we observed that SCFAs induce an increase in the level of SUMO2/3-conjugated proteins. In contrast, incubation of cells with sodium acetate, butyrate or propionate (5 mM final concentration; pH∼7.5) does not trigger any change in the SUMOylation pattern of CACO2 cells. Together, these results indicate that SCFAs, as BCFAs, modulate intestinal protein SUMOylation in a pH-dependent manner (Fig. 3C).

### BCFAs-induced hyperSUMOylation is dependent of ROS production

Butyric acid was previously reported to induce ROS (Reactive Oxygen Species) production in both IEC-6 intestinal epithelial cells and HeLa cells^10^. We thus tested whether SCFAs and BCFAs similarly induce ROS production in CACO2 and HeLa cells. For this, we used a sensitive luminol-based ROS detection assay^25^. We observed that the addition of isobutyric, isovaleric or butyric acid induce ROS production in both CACO2 and HeLa cells after 1h of incubation (Fig. 4A). This oxidative stress is transient as the level of ROS decreased between 1 and 5h of incubation (Fig. 4A). Interestingly, the oxidative stress induced by BCFAs and SCFAs is pH-dependent as no ROS were detected after incubation with sodium isobutyrate, isovalerate or butyrate (Fig. 4A). This suggests that ROS are produced only in response to the diffusion of the acidic form of BCFAs and SCFAs inside CACO2 cells.

**Figure 4:**
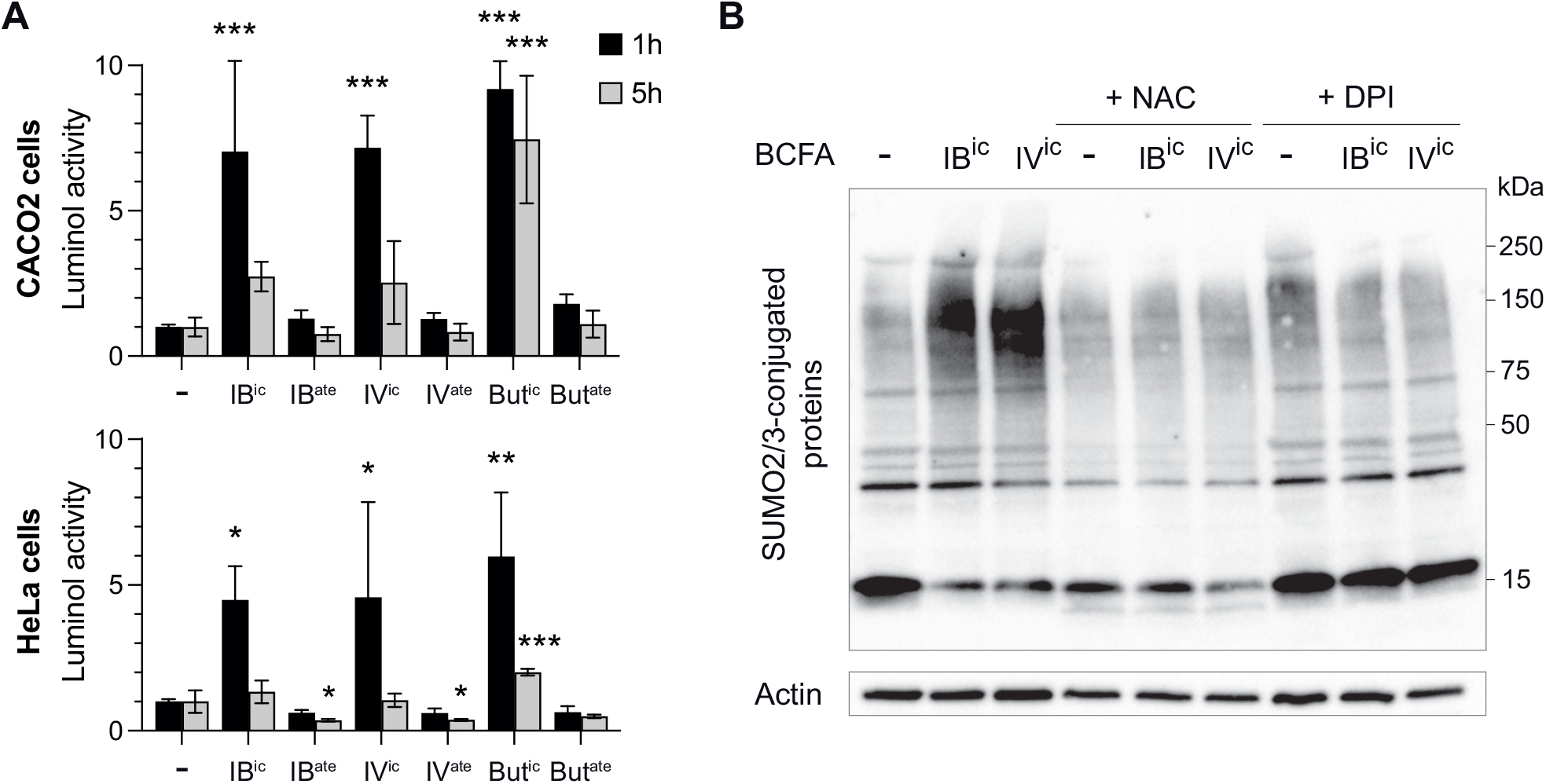
BCFAs and SCFAs induce hyperSUMOylation in intestinal cells via ROS production. A, Quantification of luminol activity in CACO2 (top) or HeLa cells (bottom) treated with isobutyric acid (IB^ic^), sodium isobutyrate (IB^ate^), isovaleric acid (IV^ic^), sodium isovalerate (IV^ate^), butyric acid (But^ic^) or sodium butyrate (But^ate^) for 1 or 5 h. Values are expressed as fold-change of untreated cells (mean ± s.d.; *n*=3; *, *P*<0.05; **, *P*<0.01; ***, *P*<0.001 versus untreated; One-way ANOVA, with Dunnett’s correction). B, Immunoblot analysis of SUMO2/3-conjugated proteins and actin levels in CACO2 cells pre-incubated or not for 30 min with 5 mM N-acetyl-cysteine (NAC) or 10 µM Diphenyleneiodonium (DPI) and then incubated for 1 h with isobutyric acid (IB^ic^) or isovaleric acid (IV^ic^).

To determine whether ROS production was responsible for the hyperSUMOylation triggered by fatty acids, we pre-incubated CACO2 cells with two ROS scavengers, N-acetyl cysteine (NAC) and Diphenyleneiodonium (DPI). These cells were then incubated with isobutyric or isovaleric acids for 5h. We observed that preincubation with oxidative stress inhibitors blocks BCFAs-induced hyperSUMOylation (Fig. 4B). Together, these results demonstrate that the acidic forms of SCFAs and BCFAs trigger the production of ROS in intestinal cells, which in turn promotes SUMO-conjugation of intestinal proteins.

### BCFAs inhibit intestinal cell deSUMOylases

Global increase in SUMOylation may result either from an increase in the SUMOylation machinery’s activity or from an inhibition of cellular deSUMOylases. As deSUMOylases were reported to be sensitive to oxidative stress^27,28^, we evaluated whether BCFAs could inhibit SUMO-deconjugation in intestinal cells. For this, CACO2 and T84 cells were incubated with isobutyric or isovaleric acids and lysed. Cell lysates were then mixed with SUMO1 or SUMO2 peptides covalently linked to AMC (7-amido-4-methylcoumarin). The activity of deSUMOylases was then quantified in these cell lysates by measuring the fluorescence intensity of AMC released by the deSUMOylase-dependent cleavage of the amide bond between AMC and SUMO (Fig. 5)^26^. We demonstrated that incubation with 5 mM isobutyric or isovaleric acids for 5 h significantly decreases SUMO deconjugation reactions in cell lysates, both for SUMO1-and SUMO2-conjugated substrates (Fig. 5). These results indicate that deSUMOylases are inhibited in response to BCFAs exposure.

**Figure 5:**
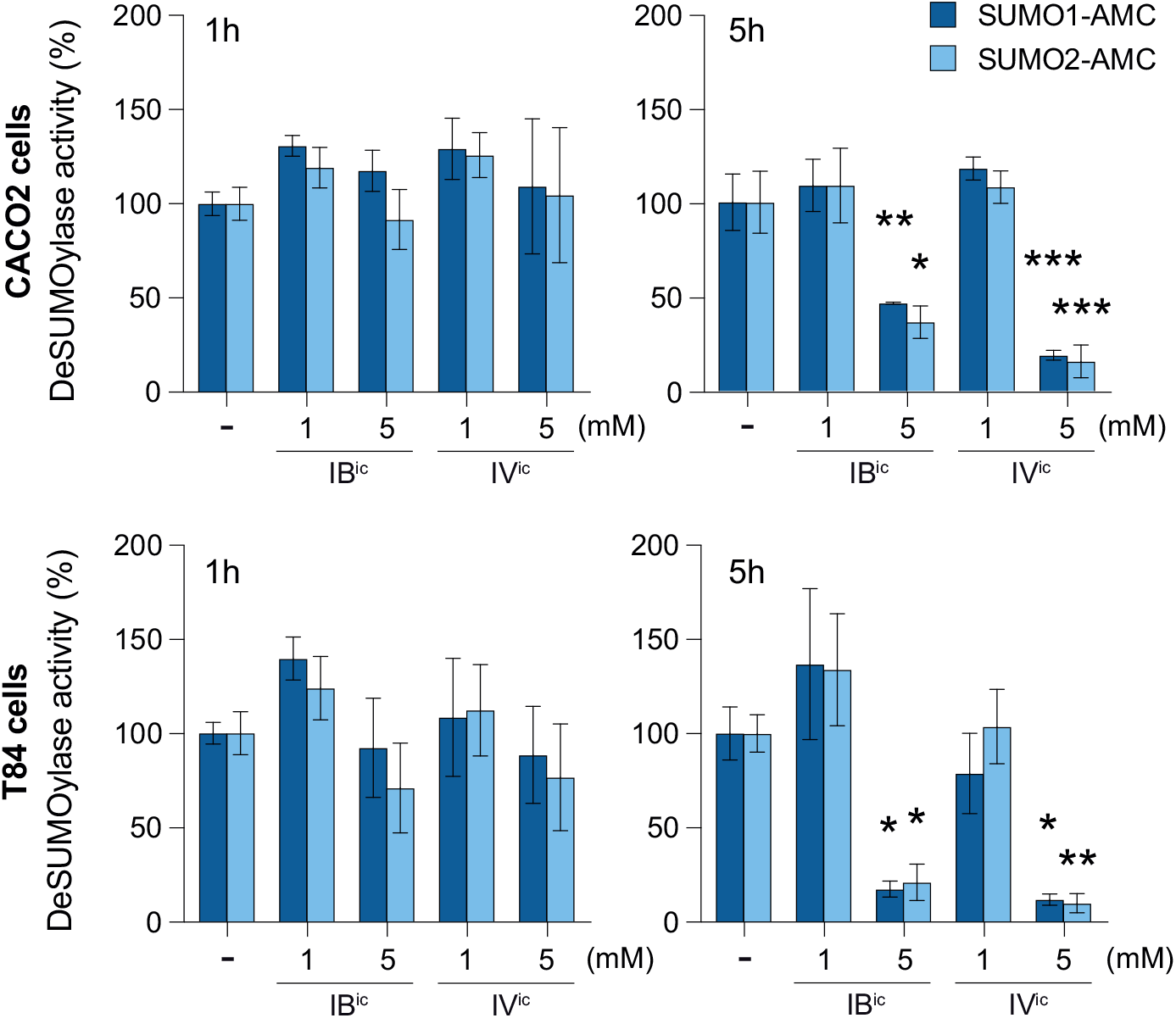
BCFAs inhibit intestinal cell deSUMOylases. DeSUMOylase activities, expressed as percentage of untreated cells, in CACO2 (top) or T84 (bottom) cells, treated or not with isobutyric acid (IB^ic^) or isovaleric acid (IV^ic^), for 1h (left) or 5h (right) (mean ± s.e.m.; *n*=4-5; *, *P*<0.05; **, *P*<0.01; ***, *P*<0.001 versus CTRL; One-way ANOVA, with Dunnett’s correction).

Of note, we quantified in parallel the expression levels of E1 and E2 SUMO enzymes in CACO2 cells treated with BCFAs using immunoblotting experiments. We observed that isobutyric and isovaleric acids do not alter the level of SAE1/SAE2 or UBC9 (Fig. S2).

Together, these results suggest that the hyperSUMOylation induced by BCFAs result from the inhibition of intestinal cell deSUMOylases.

### BCFAs-induced ROS does not affect Cullin-1 neddylation in CACO2 cells

In addition to SUMOylation, other Ubiquitin-like proteins such as NEDD8 were reported to be sensitive to oxidative stress. Previous reports established that ROS produced in response to butyric acid exposure inactivate the NEDD8-conjugating enzyme Ubc12 and trigger the loss of cullin-1 neddylation in HeLa cells^9,10^. We thus assessed whether BCFAs also decrease cullin-1 neddylation in CACO2 or HeLa cells. Interestingly, we observed that isobutyric and isovaleric acid triggers cullin-1 deneddylation after 5h of incubation in HeLa cells, similarly to butyric acid, but not in CACO2 cells (Fig. S3). This suggests that the consequences of SCFAs/BCFAs-induced ROS are cell-type dependent and that BCFAs/SCFAs do not affect neddylation in CACO2 cells.

### BCFAs promotes SUMOylation of chromatin-bound proteins

In order to decipher whether all SUMO targets are similarly affected by fatty acids-induced hyperSUMOylation, we first focused on RanGAP1, one of the main SUMO targets in human cells. We observed that the SUMOylation level of this specific target was not affected by BCFAs (Fig. S3), which suggests that only a subset of intestinal proteins are hyperSUMOylated. In order to characterize whether proteins conjugated to SUMO in response to BCFAs are located in specific cellular compartments, we performed cell fractionation assays. Very interestingly, we observed that the level of SUMO-conjugated proteins was strongly increased only in chromatin-bound fractions, thus suggesting that BCFAs promote the SUMOylation of chromatin-bound factors (Fig. 6).

**Figure 6:**
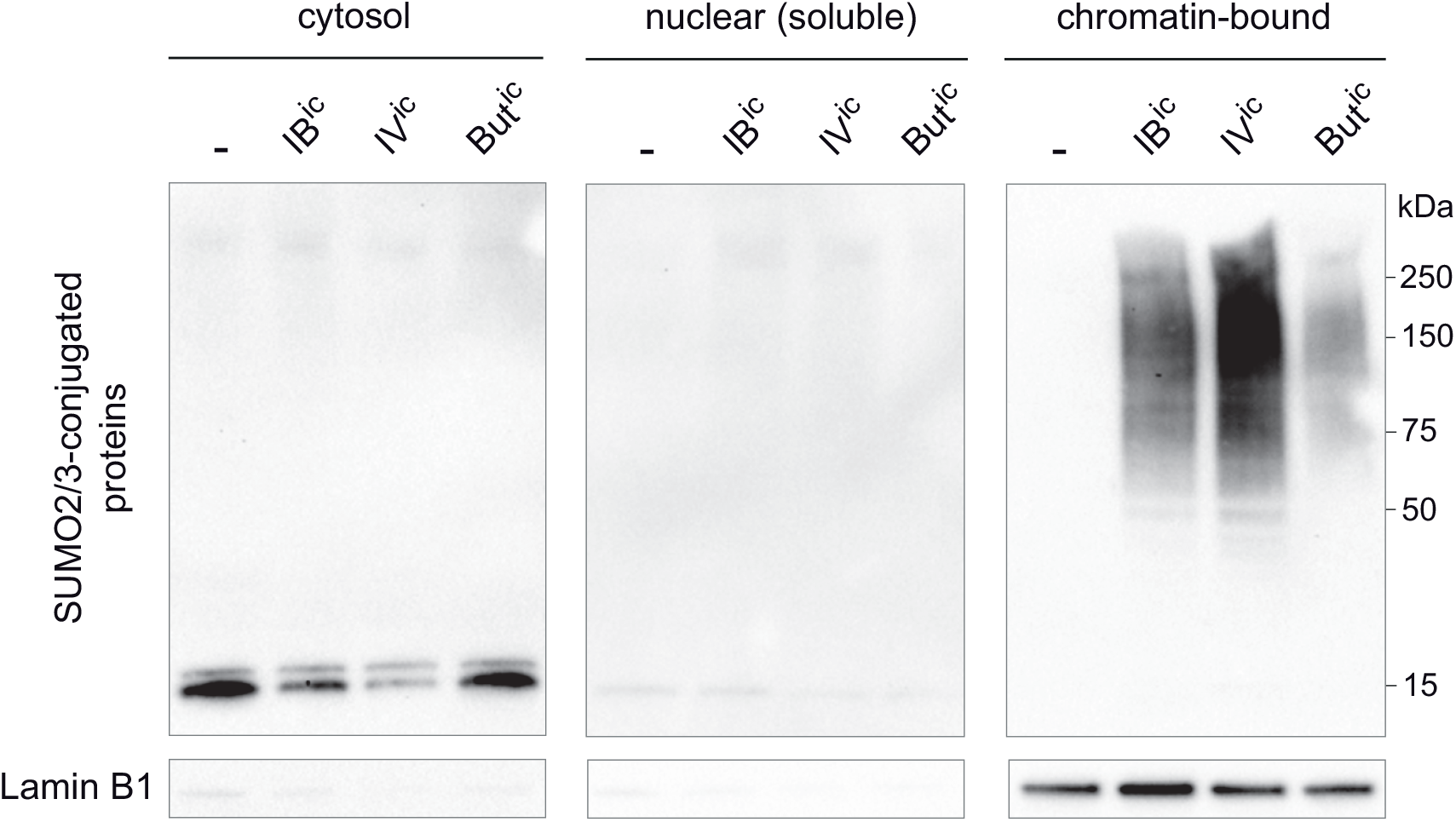
BCFAs and SCFAs trigger hyperSUMOylation of chromatin-bound proteins. Immunoblot analysis of SUMO2/3-conjugated proteins and Lamin B1 levels in cytosolic, nuclear soluble and chromatin-bound fractions of CACO2 cells incubated for 5h with isobutyric acid (IB^ic^), isovaleric acid (IV^ic^) or butyric acid (But^ic^).

### BCFAs and SCFAs-induced hyperSUMOylation impair NF-κB inflammatory responses

As SUMOylation is known to regulate inflammation^17,29^, we determined whether BCFAs/SCFAs-induced hyperSUMOylation could modulate inflammatory responses in intestinal cells. To do so, we incubated CACO2 cells with TNFα in the presence or absence of BCFAs. We then quantified the expression levels of the pro-inflammatory IL8 and CCL20 cytokines by qRT-PCR. We observed that both isobutyric and isovaleric acids downregulate the transcription of IL8 and CCL20 in response to TNFα in CACO2 cells (Fig. 7A). We then compared the respective effect of the acidic or basic forms of BCFAs and SCFAs on the expression of these cytokines. We observed that the basic form of BCFAs and SCFAs partially decrease expression of IL8 and CCL20. Interestingly, we show that the acidic forms of BCFAs and SCFAs further decrease the expression of IL8 and CCL20 to the levels of cells unstimulated by TNFα (Fig. 7A). As acidic forms of SCFAs/BCFAs trigger hyperSUMOylation in contrast to basic forms (Fig. 3), this result show that SCFAs/BCFAs-induced hyperSUMOylation dampens pro-inflammatory cytokines expression in intestinal cells.

**Figure 7:**
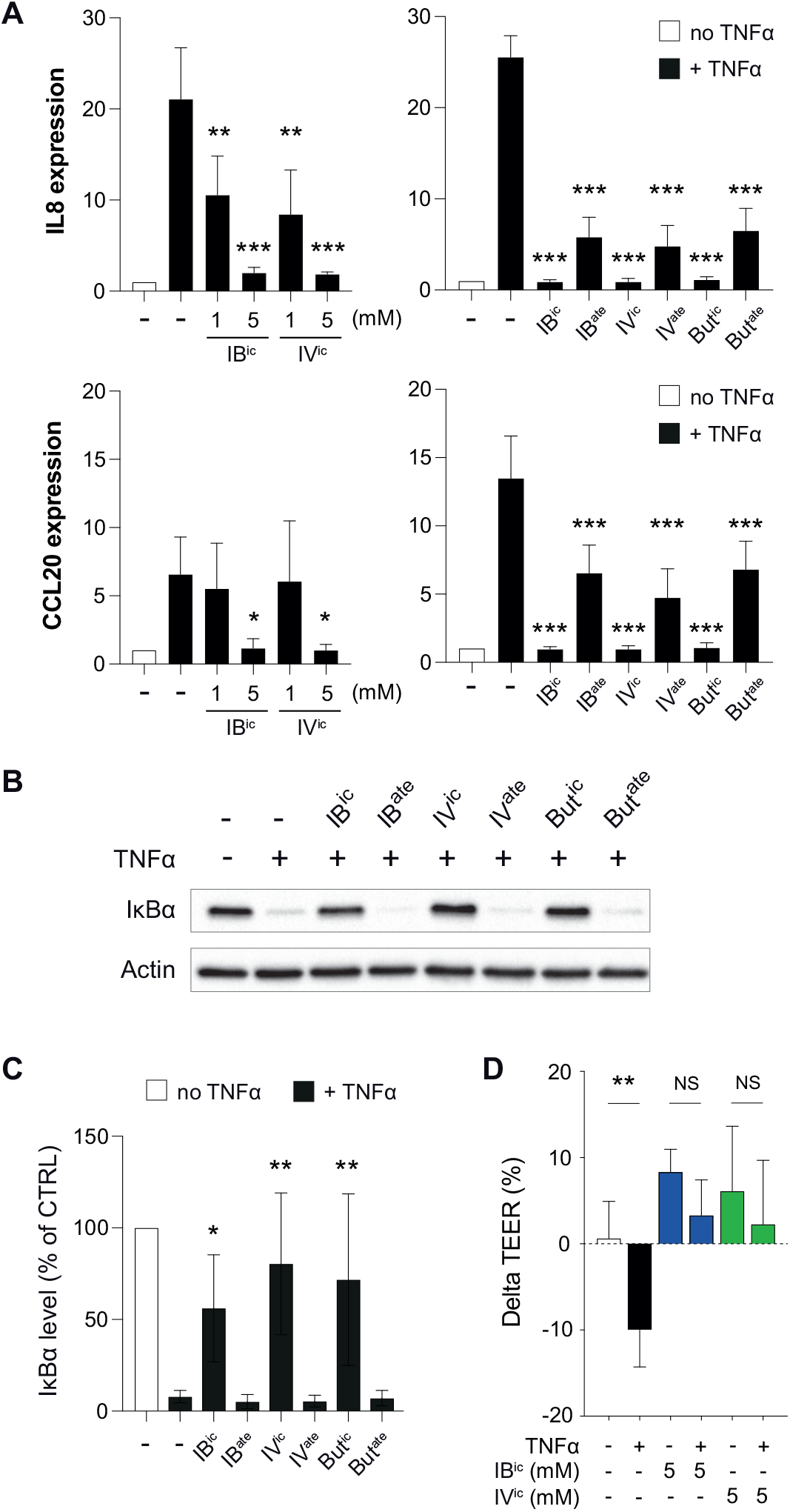
BCFAs and SCFAs dampen responses to TNFα in intestinal cells. A, Quantification of IL8 and CCL20 mRNA levels in CACO2 cells pre-treated or not for 1h with BCFAs or SCFAs and then incubated for 1h with 100 ng/mL TNFα. Values are expressed as fold change versus untreated cells (mean ± s.d.; *n*=3-4; *, *P*<0.05; **, *P*<0.01; ***, *P*<0.001 vs TNFα alone; One-way ANOVA, with Dunnett’s correction). B, Immunoblot analysis of IκBα and actin levels in CACO2 cells pre-treated or not for 1h with 5 mM BCFAs or SCFAs and then stimulated for 30 min with 100 ng/mL TNFα. C, Quantification of IκBα levels, expressed as percentage compared to untreated cells (mean ± s.d.; *n*=3; *, *P*<0.05; **, *P*<0.01 vs TNFα alone; One-way ANOVA, with Dunnett’s correction) (IB^ic^, isobutyric acid; IB^ate^, sodium isobutyrate; IV^ic^, isovaleric acid; IV^ate^, sodium isovalerate; But^ic^, butyric acid; But^ate^, sodium butyrate). D, TEER in CACO2 cells grown in Transwell, pre-treated or not for 1 hour with BCFAs and then incubated for 24 hours with 100 ng/mL TNFα. Values are expressed as TEER percent variations compared to cells before treatment with BCFAs (mean ± s.e.m.; *n*=4; **, *P*<0.01; NS, not significant; two-tailed Student’s t-test).

As IL8 and CCL20 expression is regulated by the NF-κB transcription factor, we tested whether BCFA could interfere with the NF-κB signaling pathway. To do so, we focused on the degradation of the IκBα inhibitor, which is a key step in the activation of NF-κB and a pre-requisite for NF-κB translocation into the nucleus. We quantified using immunoblotting experiments the level of IκBα in CACO2 cells incubated with TNFα in the presence or absence of BCFAs and SCFAs. We observed that isobutyric, isovaleric and butyric acids block the degradation of IκBα triggered by TNFα (Fig. 7B and 7C). This inhibition was not observed with sodium isobutyrate, isovalerate and butyrate, suggesting that the hyperSUMOylation induced by the acidic forms of SCFAs/BCFAs block IκBα degradation and thus dampen the NF-κB signaling pathway (Fig. 7B and 7C).

### BCFAs-induced hyperSUMOylation promote intestinal epithelial integrity

We finally determined whether BCFAs-induced hyperSUMOylation regulate intestinal permeability. To do so, CACO2 cells were grown for 3 weeks in Transwell systems in order to reconstitute an *in vitro* model of differentiated intestinal epithelium. Cells were then incubated with TNFα and the permeability of the obtained epithelium was monitored by measuring the transepithelial electrical resistance (TEER) between the apical and basal compartments. Treatment with TNFα induces a significant decrease in TEER after 24h, which corresponds to an increase in epithelial permeability, as previously described (Fig. 7D). Interestingly, we show that incubation of CACO2 cells with isobutyric and isovaleric acids blocks this TNFα-induced increase in epithelial permeability. This result show that BCFAs promote intestinal epithelial integrity in response to inflammatory stimuli.

## Discussion

Post-translational modifications are widely used by eukaryote cells to modulate rapidly, locally and specifically the interactions or activities of key proteins. SUMOylation plays an essential role in intestinal physiology and more particularly in epithelial integrity maintenance, by controlling cell renewal and differentiation, as well as mechanic stability of the epithelium^16,17^. Not surprisingly, several pathogens were shown to manipulate intestinal SUMOylation in order to interfere with the activity of key host factors involved in infection^18^. Most of these pathogens are decreasing SUMOylation, using independent mechanisms, which illustrates a nice example of evolutive convergence. In contrast to pathogens, the potential impact of gut commensal bacteria on SUMOylation has not been investigated. Here, we show that bacterial metabolites upregulate intestinal SUMOylation by controlling the activity of host deSUMOylases. As the SUMOylation level of a given target results from the dynamic equilibrium between conjugation and deconjugation reactions, the inactivation of deSUMOylases results in an increase in protein SUMOylation levels. Interestingly, we identified that the proteins SUMOylated in response to BCFAs/SCFAs are mainly chromatin-bound proteins. As many SUMO targets are transcription factors, we can hypothesize that BCFAs/SFAs-induced SUMOylation modifies intestinal cell gene expression^14,29^.

Our results show that BCFAs/SCFAs, by upregulating SUMOylation, dampen inflammatory responses of intestinal cells. SCFAs, and more particularly butyrate, have already been shown to have an effect on intestinal inflammation^3^. The potential effect of BCFAs on inflammation remains in contrast poorly documented. Interestingly, long-chain BCFAs (with more than 14 carbons) were shown to decrease the expression of IL8 in response to LPS in CACO2 cells and to decrease the incidence of necrotizing enterocolitis in a neonatal rat model^30,31^. Whether these effects are triggered by the acidic form of these long-chain BFCA, once translocated inside intestinal cells, remain to be determined. Of note, lactic acid, which is abundantely produced by the vaginal microbiota, also elicits anti-inflammatory responses on human cervicovaginal epithelial cells^32^. Interestingly, only lactic acid, and not lactate, prevents pro-inflammatory cytokines expression in epithelial cells, which nicely echoes our result on the anti-inflammatory properties of the acidic forms of BCFAs/SCFAs on intestinal cells^32,33^. The vaginal pH being naturally acid (pH<4), lactic acid is predominant in this environment compared to lactate. In the case of BCFAs and SCFAs produced by gut microbiota, the intraluminal pH is varying depending on the intestinal segment. This pH ranges from 5.5-7.5 in the caecum/right colon and then increases in the left colon and rectum to 6.1-7.5^34^. Eventhough the acidic forms of SCFAs/BCFAs are not predominant in these conditions, the physiological high concentrations of SCFAs/BCFAs (i.e. ∼100 mM for SCFAs) may be high enough to have a concentration of protonated fatty acids sufficient to modulate intestinal SUMOylation.

Interestingly, SUMOylation has been involved in intestinal diseases such as Inflammatory Bowel Diseases (IBD). Indeed, patients with IBD show a downregulation of the UBC9 enzyme and a decrease in SUMOylated protein levels in the colon, which correlates with disease severity^35^. These SUMO alterations, which can also be observed in a mouse model of colitis, were proposed to contribute to intestinal immune responses deregulation^35^. This hypothesis is supported by the inhibition of gut inflammation observed in response to PIAS1 E3 ligase overexpression in the intestine and the associated increase in SUMOylation^36^. Our results suggest that BCFAs/SCFAs may similarly limit inflammation in this context, by restoring SUMOylation in intestinal cells.

In conclusion, this work unveils a new mechanism used by the gut microbiota to modulate intestinal cell activies and dampen inflammation. It highlights in addition the therapeutic potential of SUMOylation targeting in the treatment of inflammatory diseases such as IBD.

## Acknowledgements

This work was supported by INSERM, Rouen University, the iXcore Foundation for Research, the Microbiome Foundation, Janssen Horizon, the European Union and Normandie Regional Council. Europe gets involved in Normandie with European Regional Development Fund (ERDF).

## Supplementary Figures

**Figure S1 : 2-methyl-butyric acid triggers hyperSUMOylation of intestinal proteins *in vitro***.

Immunoblot analysis of SUMO2/3-conjugated proteins and actin levels in CACO2 (left) and T84 (right) cells incubated with 2-methyl-butyric acid (2mBut^ic^) for 1 or 5 h.

**Figure S2 : BCFAs do not alter the expression levels of E1 and E2 SUMO enzymes**.

Immunoblot analysis of UBC9, SAE1, SAE2 and actin levels in CACO2 cells incubated for 1 or 5 h with 1 or 5 mM isobutyric acid (IB^ic^) or isovaleric acid (IV^ic^).

**Figure S3 : BCFAs do not affect Cullin-1 neddylation nor RanGAP1 SUMOylation in CACO2 cells**.

A, Immunoblot analysis of Cullin-1 and actin levels in CACO2 and HeLa cells incubated with 5 mM BCFAs or SCFAs. B, Quantification of the percentage of neddylated Cullin-1 (mean ± s.d.; *n*=3; NS, not significant; **, *P*<0.01 vs CTRL; One-way ANOVA, with Dunnett’s correction). C, Immunoblot analysis of RanGAP1 and actin levels in CACO2 cells incubated with 1 or 5 mM BCFAs. D, Quantification of the percentage of SUMOylated RanGAP1 (mean ± s.d.; *n*=3; NS, not significant; One-way ANOVA, with Dunnett’s correction) (IB^ic^, isobutyric acid; IV^ic^, isovaleric acid; But^ic^, butyric acid).

**Table S1 : Primary and secondary antibody information**.

