## Supplemental Table 1 for "Fatty acids produced by the gut microbiota dampen host inflammatory responses by modulating intestinal SUMOylation"

**Supplementary Table S1: Primary and secondary antibody information.**

| **Targeted Protein** | **Assay (dilution)** | **Species** | **Source** | **Reference** |
| --- | --- | --- | --- | --- |
| Actin | WB (1:10,000) | Mouse | Sigma-Aldrich | R5441 |
| GAPDH | WB (1:10,000) | Goat | Sigma-Aldrich | SAB2500451 |
| SUMO3 | WB (1:5,000) | Rabbit | Home-made | R205 (is2) ; {Ribet, 2017 #20} |
| UBC9 | WB (1:1,000) | Mouse | BD Biosciences | 610749 |
| SAE1 | WB (1:1,000) | Rabbit | Cell Signaling Technology | #13585 |
| SAE2 | WB (1:1,000) | Rabbit | Cell Signaling Technology | D15C11 |
| RanGAP1 | WB (1:1,000) | Rabbit | Sigma-Aldrich | R0155 |
| Cullin-1 | WB (1:1,000) | Rabbit | Cell Signaling Technology | #4995 |
| IκBα | WB (1:1,000) | Rabbit | Cell Signaling Technology | #9242 |
| Lamin B1 | WB (1:1,000) | Rabbit | Cell Signaling Technology | D4Q4Z; #12586 |
| Rabbit IgG (H+L) | WB (1:5,000) | Goat | Abliance | HRP-conjugated; BI 2407 |
| Mouse IgG (H+L) | WB (1:5,000) | Goat | Abliance | HRP-conjugated; BI 2413C |
| Goat IgG (H+L) | WB (1:5,000) | Rabbit | Dakocytomation | HRP-conjugated; P0160 |
