## Supplemental Figure 1 for "Fatty acids produced by the gut microbiota dampen host inflammatory responses by modulating intestinal SUMOylation"

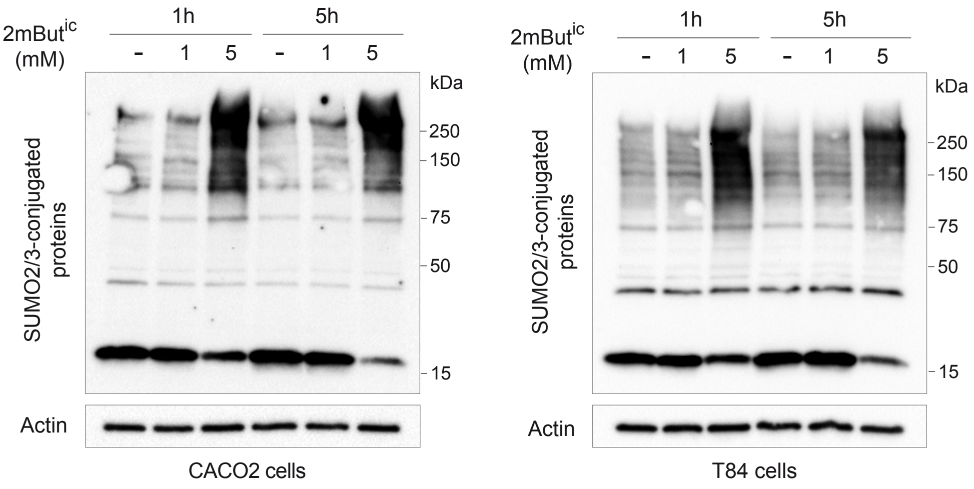


**Figure S1 : 2-methyl-butyric acid triggers hyperSUMOylation of intestinal proteins *in vitro***

Immunoblot analysis of SUMO2/3-conjugated proteins and actin levels in CACO2 (left) and T84 (right) cells incubated with 2-methyl-butyric acid ($2mBut^{ic}$) for 1 or 5 h.
