## Supplemental Figure 2 for "Fatty acids produced by the gut microbiota dampen host inflammatory responses by modulating intestinal SUMOylation"

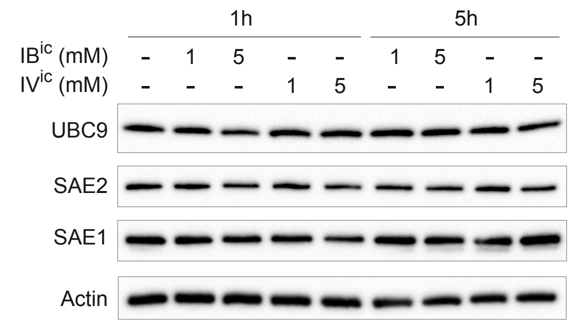


**Figure S2 : BCFAs do not alter the expression levels of E1 and E2 SUMO enzymes**

Immunoblot analysis of UBC9, SAE1, SAE2 and actin levels in CACO2 cells incubated for 1 or 5 h with 1 or 5 mM isobutyric acid ($IB^{ic}$) or isovaleric acid ($IV^{ic}$).
