## Supplemental Figure 3 for "Fatty acids produced by the gut microbiota dampen host inflammatory responses by modulating intestinal SUMOylation"

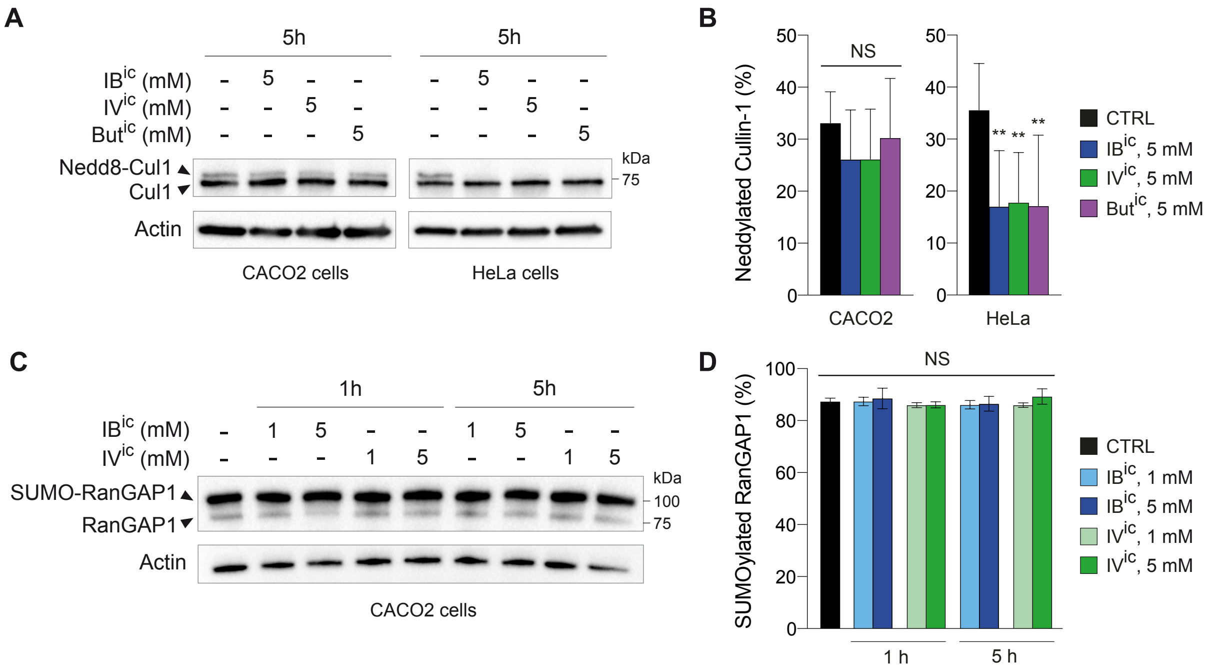


**Figure S3 : BCFAs do not affect Cullin-1 neddylation nor RanGAP1 SUMOylation in CACO2 cells**

A, Immunoblot analysis of Cullin-1 and actin levels in CACO2 and HeLa cells incubated with 5 mM BCFAs or SCFAs. B, Quantification of the percentage of neddylated Cullin-1 (mean ± s.d.; *n*=3; NS, not significant; **, *P*<0.01 vs CTRL; One-way ANOVA, with Dunnett's correction). C, Immunoblot analysis of RanGAP1 and actin levels in CACO2 cells incubated with 1 or 5 mM BCFAs. D, Quantification of the percentage of SUMOylated RanGAP1 (mean ± s.d.; *n*=3; NS, not significant; One-way ANOVA, with Dunnett's correction) ($IB^{ic}$, isobutyric acid; $IV^{ic}$, isovaleric acid; $But^{ic}$, butyric acid).
